# Summer rainfall drives adaptation with gene flow in a widespread butterfly

**DOI:** 10.64898/2025.12.12.694053

**Authors:** Lily F. Durkee, Christen M. Bossu, Kristen C. Ruegg, Brenna R. Forester, Paul A. Opler, Ruth A. Hufbauer

## Abstract

Understanding how environmental variation interacts with gene flow to shape population genomic patterns is a central goal in evolutionary biology. We investigated how geographic and environmental differences impact genomic variation in the clouded sulfur butterfly (*Colias philodice eriphyle*) by conducting whole-genome resequencing across replicated transects consisting of paired high- and low-elevation sites on both sides of a major mountain range. Despite sampling across steep environmental gradients, we found no evidence of discrete population structure, indicating high connectivity across the region. Nonetheless, significant isolation by distance – strongest in eastern populations – revealed that geographic distance still imposes limits on gene flow, and genetic diversity was also elevated in the east. Genotype-environment association analyses identified more than 16,000 loci associated with elevation, precipitation, and solar radiation. Our redundancy analysis identified precipitation as the strongest predictor of adaptive genomic differentiation, and candidate genes included those linked to melanization and thermoregulation (e.g., *TH* and *yellow*). These results demonstrate that even in a largely panmictic population, environmental variation can maintain regional-scale signals of local adaptation. Because insects are declining globally and remain underrepresented in genomic monitoring, conducting whole-genome analyses in a widespread species provides valuable context for assessing how insects today persist across such diverse landscapes and their potential for withstanding future environmental change.

## Introduction

Examining the genetic variation within and among populations is a critical first step for effective conservation management in the modern era. Natural populations can become genetically differentiated (Slatkin, 1987) or locally adapted (Kawecki & Ebert, 2004; Savolainen et al., 2013), particularly if populations are isolated (Allendorf, 1986; Mattila et al., 2012) or experience divergent selection pressures (Funk et al., 2016; Laurent et al., 2016). Genome-level analyses now enable comprehensive assessments of genetic diversity, population structure, and adaptation present across a species’ range (Allendorf, 2017; Allendorf et al., 2021; Primmer, 2009). Genomic analyses can reveal populations that may require targeted management efforts (i.e., conservation units) due to detrimental maladaptation (Brady et al., 2019) or accumulation of genetic load (Barbosa et al., 2018; Funk et al., 2012; Kardos et al., 2021). Conversely, genomic tools can reveal adaptive variants that confer fitness advantages in current or even future environments, indicating potential resilience to environmental change (Bell, 2013; Holderegger, 2015; Razgour et al., 2019; Tellier et al., 2024). Together, these tools form the foundation of conservation genomics, allowing scientists to connect genome-wide variation with population-specific evolutionary potential (Allendorf et al., 2018; Kardos et al., 2021; Whiteley et al., 2015). Yet, among animals, most genomic research has focused on vertebrates, leaving invertebrates – the majority of animal diversity – critically understudied (Bartoňová et al., 2023; Stork, 2018; Webster et al., 2023).

This gap in conservation genomic research is particularly relevant for insects, which are undergoing rapid, global declines (Harvey et al., 2022; Wagner et al., 2021; Warren et al., 2021; Webster et al., 2023). For example, the rusty-patched bumblebee, once common, has been lost from over 70% of its native range in the continental USA and is now federally listed (Colla et al., 2012; Mola et al., 2024). Recent conservation efforts for this species have focused on monitoring genetic diversity and abundance to inform recovery planning (Mola et al., 2024).

Other genetic studies of insects have focused on those that are regionally listed as endangered including the checked blue butterfly and a rare damselfly in Central Europe (e.g., Czajkowska et al., 2020; Landmann et al., 2021). However, focusing on rare species misses critical insights, as recent insect declines have been largely driven by formerly abundant taxa (van Klink et al., 2024). Studying the genomics of currently common insect species, therefore, can inform proactive conservation strategies by investigating current levels of genetic variation and the genomic basis of adaptation (Harvey et al., 2022).

The short life cycles of insects can facilitate rapid adaptation (i.e., less than 100 years) following environmental change (Carlson et al., 2014; Durkee et al., 2023; Tellier et al., 2024). This potential for rapid adaptation makes insects valuable systems for investigating how environmental variation shapes evolutionary processes (Reusch & Wood, 2007; Webster et al., 2023). In particular, elevation gradients can be used to study a species across a range of environmental conditions within relatively small spatial scales, as temperature tends to decrease with increasing elevation (Branch et al., 2017; Keller et al., 2013). Organisms whose distributions span mountain ranges are exposed to changes in environmental conditions along elevation gradients as well potential barriers to dispersal (Machado et al., 2018). Thus, a focus on such organisms can provide understanding of the combined effects of adaptation to different environments and restricted gene flow (Manel & Holderegger, 2013) by using genomic methods such as genotype-environment association (GEA) analyses that can identify signals of adaptation and associated population divergence (Capblancq & Forester, 2021; Forester et al., 2016, 2018). Conducting GEA analyses along elevation gradients allows for an assessment of adaptation to different habitats, such as those that might be later imposed due to climate change, at a single point in time (Montejo-Kovacevich et al., 2022; Peterson, 1995; Webster et al., 2023).

For our study, we focus on the Rocky Mountain subspecies of the clouded sulfur butterfly (*Colias philodice eriphyle* Edwards) (Lepidoptera: Pieridae). Clouded sulfurs are native and widespread across the US Intermountain West and western Canada, and they use a variety of legumes (family: Fabaceae) as host plants (Ezzeddine & Matter, 2008; Tabashnik, 1983). We picked this species for three reasons: First, it is a common species with few genomic assessments to date. Second, it occupies a wide range of elevations, from alfalfa fields and grasslands in the foothills of the Rocky Mountains (around 1,500m in elevation) to subalpine meadows (up to 3,000m) (Tabashnik, 1980; Watt et al., 1979), allowing for the sampling of replicated elevational gradients. Third, wing melanization (black scales) in this species increases with elevation and is heritable (Ellers & Boggs, 2002), suggesting genomic signals of adaptation are likely to be present. Melanization increases the solar absorptivity of wings, which aids in thermoregulation during flight, as butterflies will only take flight when their muscles reach a specific temperature range (30-38°C) (Ellers & Boggs, 2002, 2004; Kingsolver, 1983; Mattila, 2015). While recent genomic work has focused on hybridization of *C. p. eriphyle* with close relatives (Jahner et al., 2025), our study represents the first landscape-level genomic investigation focused solely on *C. p. eriphyle*, offering further insight on its persistence across diverse habitats and establishing a baseline for detecting future change.

We used a conservation genomic approach to investigate genetic diversity, population structure, and adaptation of *C. p. eriphyle* across elevation transects positioned east and west of the North American continental divide. Each transect consisted of a paired “high-elevation” site in the subalpine and a “low-elevation” site in the foothills. We conducted whole genome resequencing on 165 individuals across 15 sites in Colorado and Wyoming, USA, and conducted a genotype-environment association analysis to explore the potential genomic basis of adaptation across space and altitude. We used these methods to address the following questions: (1) How do differences in environmental conditions across replicate Rocky Mountain elevational transects influence a) genetic diversity, b) population structure, and c) signals of adaptation between subalpine and foothills habitats? (2) What genes underlie adaptation, and how do they relate to known morphological variation in this species? Based on the ecology of *C. p. eriphyle* (described by Watt et al., 1979) and our sampling design, we formulated the following expectations: Given that *C. p. eriphyle* is regionally abundant, we anticipated relatively high genetic diversity across the sampling region. Additionally, because our study covered a relatively small geographic area (∼10,000 km^2^) and populations are often continuous, we predicted limited population structure overall due to gene flow among sites. Nevertheless, the highest elevations of the Rocky Mountains lie outside the species’ elevational range and could act as a barrier to dispersal, in which case we would expect population structure to be aligned with the continental divide, with distinct eastern and western populations. Our sampling design also spanned a wide range of environmental conditions, particularly between high- and low-elevation sites, which we predicted would be subject to divergent natural selection and exhibit clear genomic signals of adaptation. Finally, we expected the GEA to identify loci associated with environmental variability and genes linked to adaptation to colder temperatures (e.g., wing melanization). By evaluating these predictions, we aimed to better understand how elevational gradients and potential dispersal barriers shape population genomics in a widespread insect species.

## Methods

### Field collection

In June through September 2022, we collected *Colias p. eriphyle* butterflies from replicated elevational transects consisting of seven low- and eight high-elevation sites on both sides of the continental divide in Colorado and southern Wyoming, USA (Figure 1). Low-elevation sites were all less than 2,000m in elevation and consisted of farmland, neighborhood parks, or meadows containing alfalfa or clover. High-elevation sites were all located in subalpine meadows with an average elevation of ∼2,500m. Sites were paired geographically, with one high and one low site identified that were geographically closer together than they were to other sites (other than pairs E1, E2, and X; Figure 1b). The northernmost site (*Wy*) did not have a low-elevation pair due to geographic constraints in the surrounding area. All high-low site pairs had a difference in elevation of at least 1,000m. We collected 6-20 individuals per site with mesh butterfly nets, and a GPS coordinate was taken for every individual (except for individuals collected at *Wy*). Site area was determined using these GPS coordinates and the *areaPolygon* function from the *geosphere* package in R (Hijmans, 2022). We assessed the relative abundance of *Colias* species at each site using Pollard Walks (a standard measure of relative abundance of butterflies) (Pellet et al., 2012; Pollard, 1977), and measured temperature and wind speed using a Kestrel device. Collections were only made when conditions for that day included an air temperature greater than 20°C and windspeed less than 20 kph, and when relative abundance was greater than 1.

**FIGURE 1.**
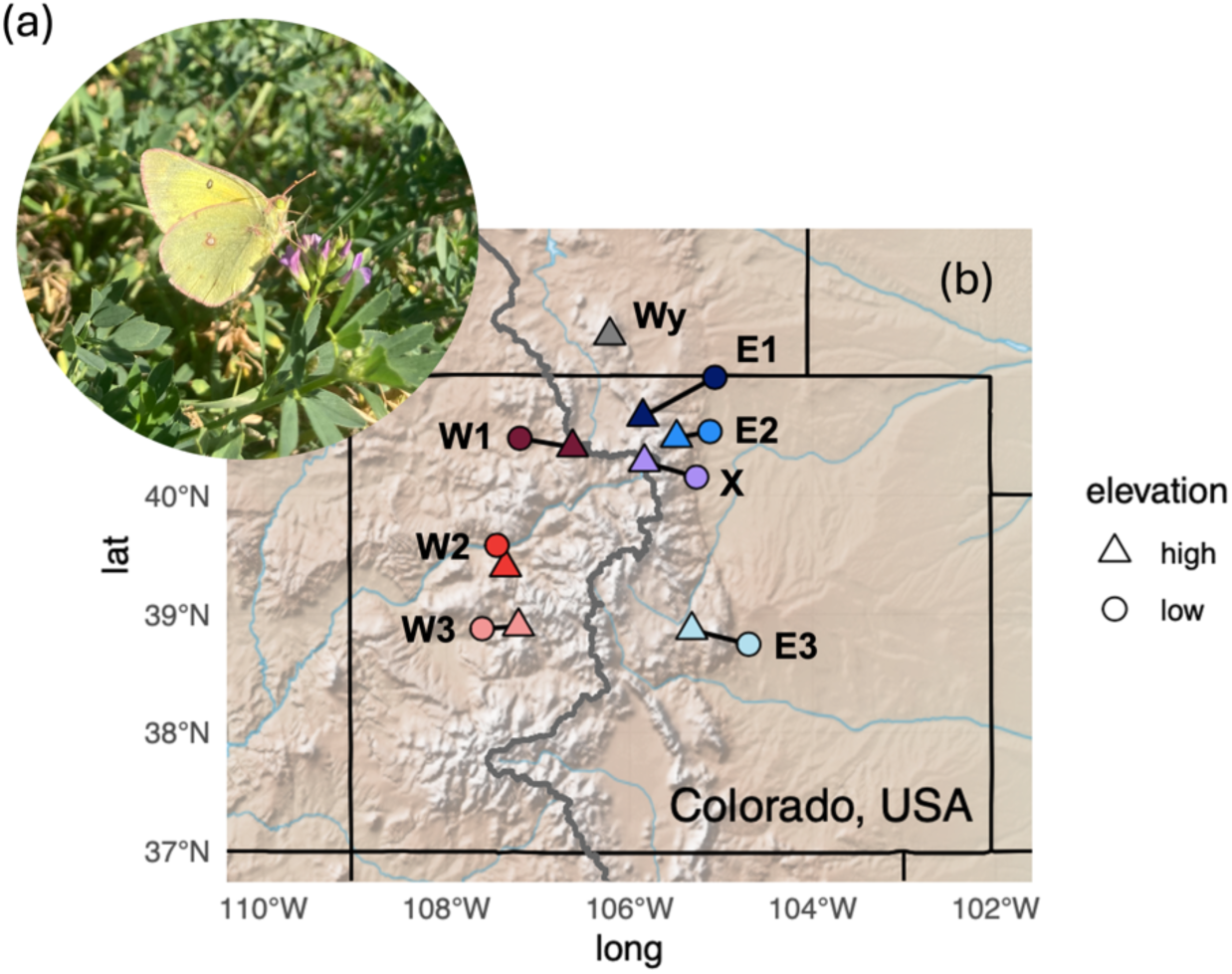
**(a)** Photo of a male *Colias p. eriphyle,* taken in Loveland, Colorado by LF Durkee. **(b)** Map of all sites sampled for *C. p. eriphyle* in Colorado and southern Wyoming. Transects connect paired high and low sites (triangles and circles, respectively) both east (E; blue shades) and west (W; red shades) of the continental divide (gray line). The X site pair (purple points) goes “across” the continental divide. The northernmost site, *Wy* (gray triangle), does not have a paired site due to geographic constraints.

Netted individuals were placed in glassine envelopes and transported to Colorado State University and were frozen alive or within 48hr of death at −80°C. Permission to collect individuals was obtained from the relevant agency (U.S. National Park Service permit no. ROMO-00257, U.S. Forest Service, City of Fort Collins, or private landowners) prior to the field season.

### Sample preparation and resequencing

We extracted DNA from the thorax of up to 20 individuals per site using Qiagen DNeasy Blood and Tissue Kits. We assessed DNA yield using *qubit* assays and evaluated sample quality by running each on agarose gels. Library preparation was completed following Schweizer & DeSaix (2023) on up to 12 of the highest quality samples per site. In total, we sequenced 170 *C. p. eriphyle* individuals from 15 collection sites (6-12 individuals per site; Table 1) on three lanes of Novogene HiSeq 4000. Additionally, we included a related species, *Colias eurytheme,* the orange sulfur, which is known to hybridize with *C. p. eriphyle* (Dwyer et al., 2015) and can be difficult to distinguish visually. We included six individuals of this species to assess hybridization or potential misidentification of our clouded sulfur individuals.

**TABLE 1.**
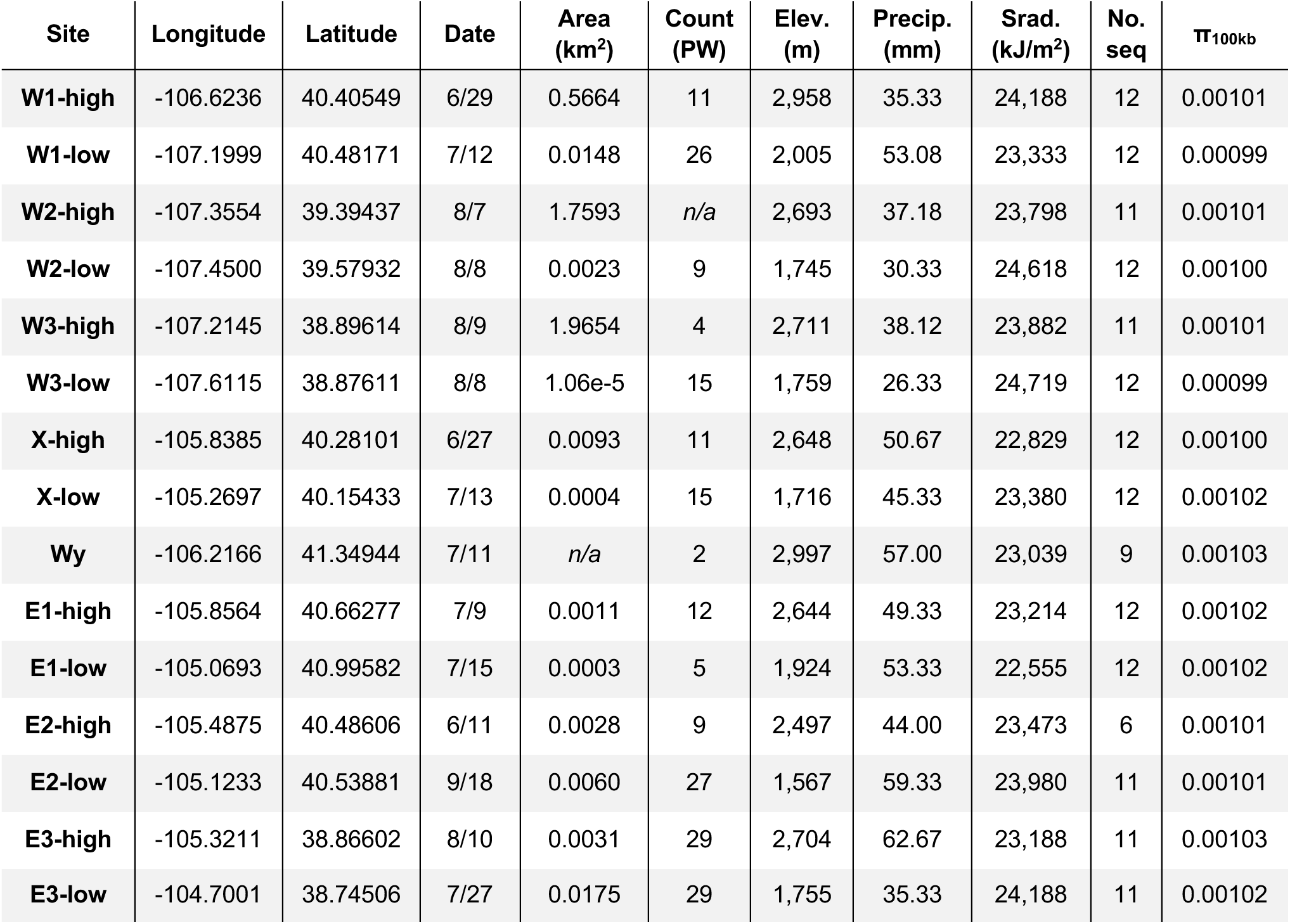
Collection site information, including site names, locations, collection date (month/day in 2022), area (km^2^), Pollard Walk (PW) count, average elevation (elev, m) from *elevatr,* as well as average monthly precipitation (precip, mm) and average daily solar radiation (srad, kJ/m^2^) for the summer months (June-August). Sample and diversity measures are also shown, including the number of butterflies from each site that were included in analyses (no. seq) along with the mean of window-based π (100kb window).

### Bioinformatic processing

All bioinformatic processing and analyses were completed using the RMACC Alpine computing cluster managed by Colorado State University and University of Colorado, Boulder. Raw FASTQ files were trimmed to remove low-quality paired-end sequences using *TRIMMOMATIC* (Bolger et al., 2014; Lou & Therkildsen, 2022). We mapped the trimmed FASTQ files to the annotated clouded yellow butterfly (*Colias crocea*) reference genome (Ebdon et al., 2022) (European Nucleotide Archive, PRJEB42949) using *BWA* (Li & Durbin, 2009), added read groups using *Picard* (Broad Institute, 2019), and removed PCR duplicates with *samtools* (Danecek et al., 2021). Individual coverage on average was 5.0x, with a wide range of read depth (1.3-14.7x) (Figure S1). To remove potential coverage effects that can mask geographic signals, we downsampled the bam files using *samtools* to 4x coverage (Lou & Therkildsen, 2022). Three closely related individuals (based on the proportion of alleles identical by descent) were identified using *NGSRelate* (Korneliussen & Moltke, 2015), and removed from downstream analyses, as related individuals can skew assessments of population structure (Coleman et al., 2016). Exploratory analyses were then completed using genotype likelihoods in *ANGSD* (Korneliussen et al., 2014) using *pcangsd* to create principal component analyses (Figure S2). One individual was removed because it was misidentified; another was removed due to suspected hybridization with *C. euytheme*.

With the remaining 165 individuals, we proceeded with a *GATK* pipeline and used the *ANGSD* analysis as a check of the validity of subsequent analyses with called genotypes. We processed bam files to call genotypes using GATK4 *HaplotypeCaller* to create GVCF files with mean *QUAL* score >20 (Miller et al., 2023; Poplin et al., 2018) using a minor allele frequency filter of 5-95% (Hemstrom et al., 2024; Lou et al., 2021). We created genomic databases using *GenomicsDBImport*, genotyped the GVCFs using *GenotypeGVCFs,* and removed systematic errors from the VCF file using *VariantFiltration* and *SelectVariants* (McKenna et al., 2010; Poplin et al., 2018) to allow only biallelic sites, <80% missing data per site, and a *QUAL* score >30 (Auwera et al., 2013; Lou et al., 2021). *NA* values were marked as missing using *bcftools* (Danecek et al., 2021). Imputation was completed using *beagle v4.1* (Browning & Browning, 2016) when required for downstream analyses (Forester et al., 2018). Lastly, we conducted linkage disequilibrium (LD) pruning using *plink* (Purcell et al., 2007) to retain independent loci (*R^2^* > 0.1 within 50-SNP sliding window) recommended for analyses with *ADMIXTURE* (Alexander et al., 2009).

### Assessment of genetic diversity

The amount of genetic diversity present in a population impacts its ability to persist and adapt to future challenges (Kardos et al., 2021; Mastretta-Yanes et al., 2024). We approximated this key metric using window-based nucleotide diversity (π), calculated using *vcftools* (Danecek et al., 2011) with a window size of 100kb (Martin et al., 2016; Talla et al., 2019).

### Population structure and gene flow

Exploratory analyses of population structure were completed using principal component analyses (PCA) with the R package *vegan* (Oksanen et al., 2024). We then evaluated the generated PCAs using *F-*statistics generated by multivariate analysis of variance (MANOVA) tests using the principal components (PC1 and PC2) as response variables and site-specific variables as predictors (e.g., [PC1, PC2] ∼ site) similar to Flanagan et al. (2016). We identified principal components explaining significant levels of genetic structure using broken stick criterion (Forester et al., 2018; Jackson, 1993). We then used *F_ST_* to assess differentiation among groups.

We tested for isolation by distance by evaluating the correlations between pairwise genetic distances and geographic distance. To do this, we calculated pairwise *F_ST_* using *plink* and estimated geographic distance between each population (using the *geosphere* package in R, which provides Haversine values, in units of meters) (Hijmans, 2022; Mikheyev et al., 2013; Trense et al., 2021). Slatkin’s D was calculated as 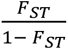 (Slatkin, 1991). We also completed a Mantel test to evaluate the correlation between genetic distance and geographic distance matrices using the “Pearson” method and 999 permutations using *vegan* (Cheek et al., 2022; Manel & Holderegger, 2013). Lastly, we evaluated ancestry using *ADMIXTURE* to test for the number of ancestral populations (*K*) that best fit the data using 5-fold cross-validation (i.e., five validation runs were performed and then averaged over). We evaluated *K*=1-10 using on the LD-pruned VCF file in *plink* format, and we chose the value for *K* with the lowest average prediction error (Alexander et al., 2009).

### Adaptation: Genotype-environment association analysis

To identify genetic variation correlated with environmental and climate variables, we used a redundancy analysis (RDA), a multivariate genotype-environment association (GEA) method that identifies genomic signatures of selection related to environmental variables (Capblancq & Forester, 2021; Forester et al., 2018). The individuals from population *Wy* (*N* = 9) were excluded from this analysis because individual GPS coordinates were not taken in the field. For all other individuals (*N* = 156), average monthly precipitation (mm), maximum temperature (°C), daily solar radiation (kJ/m^2^), and wind speed (m/s) data were extracted for each individual’s GPS coordinate from *WorldClim* raster files for monthly values at 30s (∼1km^2^) spatial resolution from 1970-2000 (Fick & Hijmans, 2017). Most of our collection sites were smaller than 1km^2^, so individuals within sites generally had either the same or very similar climatic variables. We then took the mean values for the summer months (June-August), because this is when most of the collection was completed (13 of 14 sites; Table 1) and when *Colias* flights are most likely to occur (Watt et al., 1979). Elevation (in meters) associated with each individual was obtained using the *elevatr* package in R for a greater resolution than provided by *WorldClim* (Hollister et al., 2023). To avoid including collinear predictors, we removed highly correlated (Pearson’s *r* > 0.8) climate variables (Dormann et al., 2013) resulting in three retained predictors: elevation, total monthly summer precipitation (averaged over June-August), and daily summer solar radiation (hereafter, “precipitation” and “solar radiation”) (Figure S3). We chose precipitation and solar radiation over wind speed because precipitation has been shown to shape insect distributions (Kellermann & van Heerwaarden, 2019) and solar radiation is important for thermoregulation in *Colias* butterflies (Kingsolver & Buckley, 2017). We retained elevation rather than temperature to account for other environmental shifts that occur along elevation gradients, for example oxygen levels, air pressure, and snowpack, as these variables are known to be correlated with elevation (Montejo-Kovacevich et al., 2022; Rebetez, 1996).

We identified SNPs associated with the multivariate environment by identifying those with constrained ordination axis loadings in the tails of the distribution using a standard deviation cutoff value of 3 (equivalent to *p* < 0.0027) along one or more of the RDA axes (Forester et al., 2018). To identify genes associated with each candidate loci, we used the *closest* function in *bedtools* (Quinlan & Hall, 2010) to intersect the SNP positions with the annotation of the reference genome, and kept loci found within 25kb of each gene, an estimated distance before linkage disequilibrium breaks down (Backström et al., 2006). We further analyzed the function of identified SNPs using the program *SnpEff* (Cingolani et al., 2012), which predicts the effects of genetic variants on gene function by classifying them by their impact (low, moderate, high, or intergenic/modifier) on protein-coding regions and regulatory genomic elements.

### Discovering genes underlying adaptation

Finally, instead of investigating the function of each annotated gene or gene family (Kellermann & van Heerwaarden, 2019), we searched for genes involved in three pathways critical for adaptation to environmental change in insects: (1) Wing melanization (Zhang et al., 2017), due to the association of melanin with adaptation to elevation in this butterfly (Ellers & Boggs, 2004), (2) heat shock protein synthesis, which is well-documented to aid in thermal stress tolerance in insects (Banfi et al., 2025), and (3) cytochromes P450, which are studied primarily in pest insects and underly adaptation to host plant toxins and other environmental challenges (Calla et al., 2017; Scott et al., 1998). We then examined the minor allele frequency (MAF) at each locus to identify correlations with our environmental variables using the *freq* function in *plink* (Purcell et al., 2007). Individuals were pooled by site, and significant (*p* < 0.05) correlations between MAF and elevation, precipitation, and solar radiation were identified using simple linear models in R. Thus, the candidate genes we identify meet the following criteria: (1) association with significant SNPs identified in the RDA, (2) documented involvement with one of the three pathways discussed above, and (3) statistically significant correlations between MAF at the locus and one or more of our environmental variables.

## Results

### Summary of sampled individuals

Butterflies were collected from 15 sites in a paired design throughout Colorado and southern Wyoming (Figure 1). The highest elevation site (*Wy*) was located in Wyoming’s Snowy Range; the highest eastern site in Colorado (E3-high) was from an outdoor learning center near Florrisent and the highest western site (W1-high) was located on Rabbit Ears Pass. Sites ranged in area from 10m^2^ (a small family farm near Paonia, CO) to 2km^2^ (subalpine meadow on Kepler Pass near Crested Butte, CO) (Figure 1; Table 1). In general, low-elevation sites were both smaller in area and had higher relative abundance of *Colias* (compare white vs. grey rows in Table 1), which is consistent with past studies (Tabashnik 1980, 1983) that observed high densities and low dispersal of *C. p. eriphyle* on alfalfa (*Medicago sativa*), a host plant present at six of seven low-elevation sites. The high-elevation sites mostly spanned a larger area and had sparser host plant availability of native legumes, such as clover (*Trofolium*) or vetch (*Vicia*).

Bioinformatic processing and variant detection with GATK4 resulted in 165 *Colias p. eriphyle* individuals being included in our downstream analyses, comprised of 1,454,624 SNPs and 31 scaffolds from the alignment with the reference genome of sister species *Colias crocea.* Scaffolds 1-30 represent autosomal chromosomes, followed by the sex chromosome *Z*.

### Assessment of genetic diversity

Nucleotide diversity (π) was around 0.0010 across sites, which is comparable to estimates for other species of butterflies in the Pieridae family, which found genome-wide π estimates to range from 0.001-0.004 (Talla et al., 2019; Zheng et al., 2023). Notably, nucleotide diversity was higher in the east (average π = 0.001075) compared to the west (π = 0.001058) of the continental divide (t-test: *t* = 4.332, *p* < 0.001), and this trend was linear with respect to longitude (t-test: *t* = 2.658, *p* = 0.02) (Figure 2a; Table 1).

**FIGURE 2.**
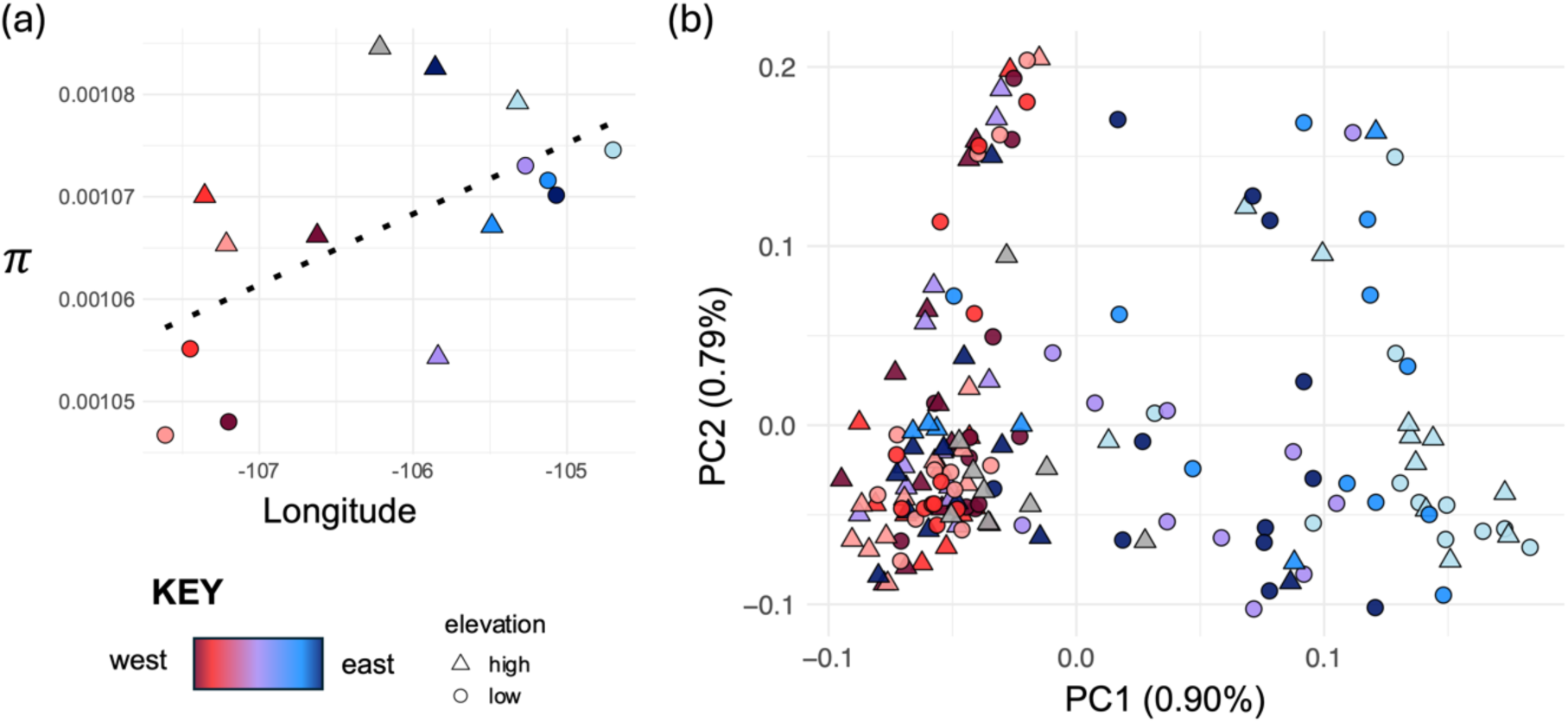
**(a)** Nucleotide diversity (π) over longitude, indicating that there is greater diversity in the east (blue shaded points) compared to the west (red shaded points). The positive, linear trend is shown with the dotted line. **(b)** Principal component analysis of all individuals (*N* = 165) across all sites (*N* = 15). Each point represents an individual butterfly: red shades represent butterflies from the west, blue shades represent butterflies from the east, triangles represent high-elevation sites, and circles represent low-elevation sites. The colors and shapes match the scheme in the map of the collection sites (Figure 1). Note that neither axis explains substantial variation (0.90 and 0.79%). Points appear to cluster more strongly by color, which corresponds to their position east vs. west of the divide. Triangles and circles show no consistent clustering, signaling weak differentiation of high and low elevation sites.

### Population structure and gene flow

We evaluated population structure and the possibility for restricted gene flow using a variety of methods. Overall, the explanatory power of the first PC axis was less than 1% (Figure 2b), and none of the PC axes exceeded the broken-stick criterion when visualized with a scree plot (Figure S4; Forester et al., 2018). The multivariate analysis of variance (MANOVA) suggested that collection site was a significant predictor of PC1 and 2 (MANOVA: *F_14,150_* =7.982, *p* < 0.001), which suggests that though the population structure among collection sites was weak, it was still greater than expected by chance. Additionally, we used *F*-statistics generated by the MANOVA and weighted *F_ST_* to assess differentiation between groups of sites. For the east vs. west sites, the MANOVA indicated strong differentiation (*F_1,163_* = 74.63, *p* < 0.0001), and weighted, global *F_ST_* = 0.0017. For high vs. low sites, the MANOVA indicated weaker differentiation (*F_1,163_* = 9.62, *p* = 0.0001) and the weighted *F_ST_* was also smaller (0.00045). Both *F_ST_* values were close to zero, indicating that population structure was low between east-west and high-low sites, however the significant MANOVA tests suggest that there is some restricted gene flow between these groups. Additionally, the *F-*statistic and weighted *F_ST_* values for the east-west comparison were 8X and 4X greater than the high-low comparison, respectively, suggesting differentiation was driven more by restricted gene flow over the continental divide than between high and low sites.

Second, we used *ADMIXTURE* (Alexander et al., 2009) to infer the number of populations represented in our 15 collection sites. This analysis revealed that the best-supported number of ancestral populations (*K*) by cross-validation (CV) was *K* = 1. This suggests that the butterflies in our study likely descend from a single ancestral population and represent a single genetic cluster, providing additional evidence for high connectivity among individuals at the collection sites.

Lastly, we evaluated isolation by distance (IBD) using a Mantel test to evaluate whether sites farther apart were more genetically differentiated than sites nearby (Figure 3). The analysis showed that the two distance matrices, Slatkin’s D (distance in *F_ST_*) and Haversine (geographic distance), were significantly correlated (*r* = 0.4692, *p* = 0.001), indicating a robust pattern of isolation by distance among sites. Furthermore, we found a stronger signal of IBD in eastern sites (*r* = 0.600, *p* = 0.011) compared to western sites (*r* = 0.3914, *p* = 0.032) (Figure S5).

**FIGURE 3.**
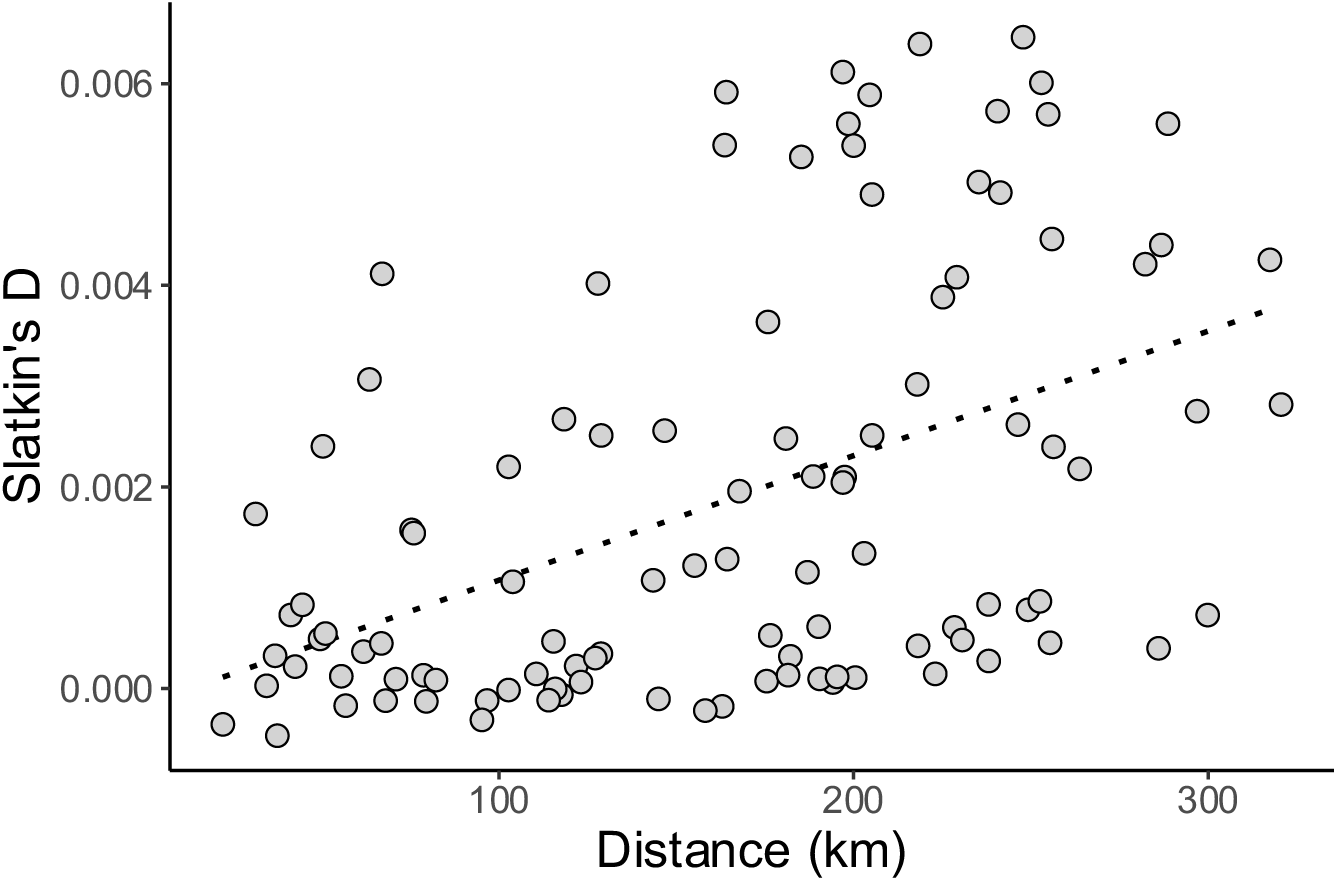
Distance among sites (Haversine distance, km) is positively correlated with Slatkin’s D, a measure of genetic distance 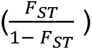. The dashed line shows the fitted linear regression model.

### Adaptation: Redundancy analysis

We investigated loci associated with the environment using a redundancy analysis (RDA), which revealed signals of differentiation due to genomic associations with environmental variables, or hereafter, adaptive divergence. No correction for population structure was included because none of the axes in the PCA exceeded the broken-stick criterion (Figure S4), as liberal inclusion of spatial correction can decrease power to detect loci under selection (Forester et al., 2018). The RDA indicated that elevation, precipitation, and solar radiation were significantly associated with genetic variation (*F_3, 31005_* = 1.107; *p* < 0.001). Examination of RDA model coefficients showed that precipitation loaded strongly on both RDA1 (loading value = 0.0081) and RDA2 (loading = −0.0082), suggesting that precipitation explains the greatest proportion of adaptive genomic variation across sites. Elevation and solar radiation exhibited weaker loadings on RDA1 (−0.00013 and 0.000025, respectively), and both loaded strongest on RDA2 (−0.00013 and −0.00018. respectively). On RDA3, the coefficients of precipitation, elevation, and solar radiation along this axis were comparatively small (−0.0013, 0.00005, −0.00012, respectively), indicating that RDA1 and RDA2 capture the primary environmental signals. Together, these results identify precipitation (or, environmental variation strongly correlated with precipitation) as the primary environmental driver of adaptive divergence among individuals.

As expected, the distribution of butterflies in the ordination plots shows that individuals (circles or triangles) were arranged in the ordination space relative to their relationship with the environmental predictors (purple arrows, Figure 4). We observe three patterns in the RDA consistent with adaptation. First, patterns associated with elevation revealed clear genomic separation between high- and low-elevation individuals that was not apparent in the PCA, indicating potential adaptive divergence associated with altitude and related environmental factors (Figure 4a–b, triangles). Second, individuals from paired high and low sites do not cluster together despite being separated by 45km on average, suggesting stronger associations with environment rather than geographic proximity. Finally, individuals within sites clustered most distinctly by position (east vs. west) relative to the continental divide, a pattern primarily driven by variation in precipitation (Figure 4a; note the angle of the precipitation axis relative to the dashed line).

**FIGURE 4.**
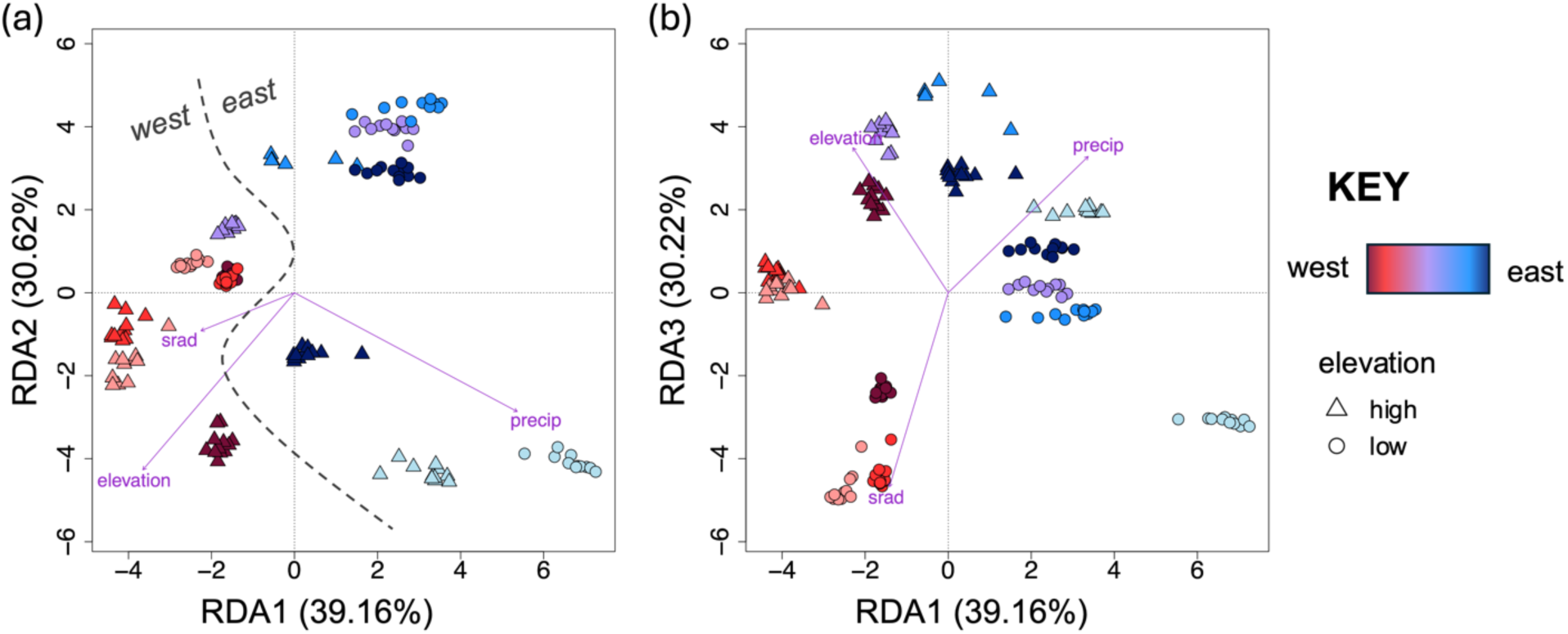
Redundancy analysis (RDA) ordination plots showing environmentally-associated genetic divergence. **(a)** Shows the plot of RDA2 vs. RDA1, and **(b)** shows RDA3 vs. RDA1. Each point represents an individual butterfly: red shades represent butterflies from the west, blue shades represent butterflies from the east, triangles represent high-elevation sites, and circles represent low-elevation sites (Figure 1). Each purple vector represents the environmental predictors. The arrangement of the points represents their relationship with the ordination axes, which are linear combinations of the environmental variables. The dashed line separates eastern from western populations.

### Loci and genes underlying adaptation

In total, we identified 16,492 significant loci, 51.4% associated most strongly with precipitation, 29.5% associated most strongly with elevation and 18.5% associated most strongly with solar radiation. The association of a majority of these loci with precipitation highlights the importance of precipitation for adaptation in this species. Of these candidate loci, 15,630 were located within 25kb of 4,082 named genes. In terms of gene function, 762 of the total loci (4.65%) were nonsynonymous variants and were predicted to have moderate effect on downstream protein function, 24.5% had low effect (synonymous and splice region variants), and 70.8% were identified as intergenic or modifiers. These results suggest that while most adaptive signals occur in noncoding regions, a subset of protein-altering variants may contribute directly to genomic adaptation.

From these loci, we identified genes meeting two criteria: functional involvement in key adaptive pathways (wing melanization, heat shock protein synthesis, or cytochrome P450 activity), and significant correlations between allele frequencies and environmental variables. Four genes or gene families – *TH, yellow, hsp70*, and *CYP6B6* – met our criteria as candidates. Regression analyses showed that these loci were associated significantly with changes in summer rainfall (Figure 5a), and they had no significant correlation with elevation or solar radiation. These genes are located across the genome (chromosomes 5, 6, 19, and *Z*; Figure 5b). We suggest that future studies explore the potential biological function or adaptive significance of these candidate genes, particularly with respect to precipitation for this species (Montejo-Kovacevich et al., 2022; Higgins et al., 2015).

**FIGURE 5.**
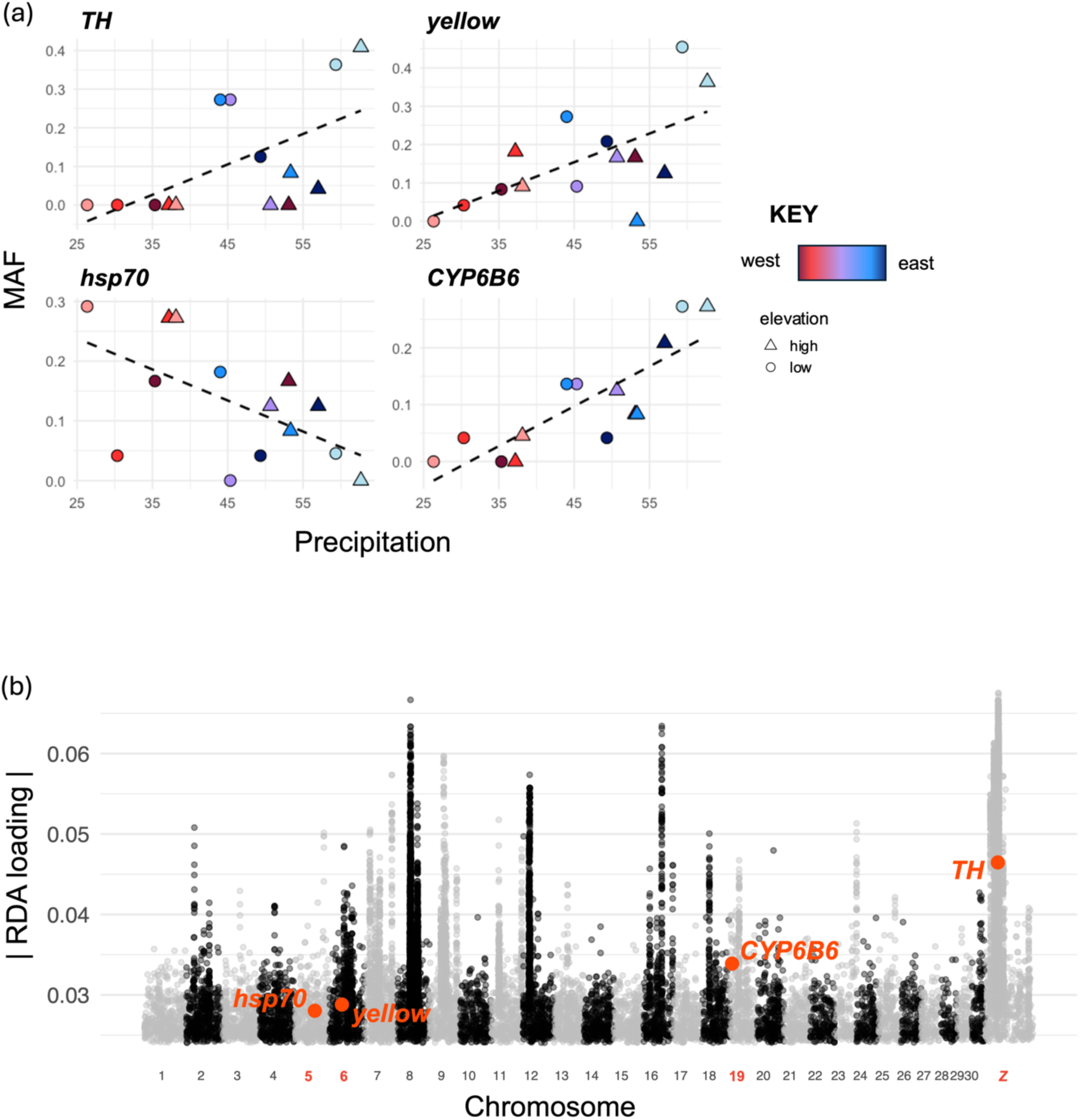
Four SNPs located within the coding regions of *TH, yellow, hsp70,* and *CYP6B6* genes or gene families, respectively, met our criteria for gene discovery. Shown are **(a)** the statistically significant, linear association of minor allele frequency (MAF) with precipitation (average summer monthly rainfall, measured in mm), and **(b)** a Manhattan plot showing the location of each SNP along the genome, with the absolute value of the RDA loading values on the y-axis and the chromosome number, labeled 1-30 and *Z*, on the x-axis.

## Discussion

We conducted whole genome sequencing to evaluate how elevation (foothills vs. subalpine) and geographic position (east vs. west of the Rocky Mountains) affects population genetic diversity and structure in clouded sulfur butterflies (*Colias p. eriphyle*), and we evaluated evidence for adaptation using a genotype-environment analysis. We predicted the following: (1) genetic diversity would be high, (2) population structure would either be panmictic or be split east-west of the continental divide, and (3) the GEA would reveal genomic signals of adaptation to elevation and environmentally-associated loci and genes. Evaluating genomic population structure on a regional scale is important for understanding more localized patterns of population connectivity and adaptation, allowing us to identify vulnerable subpopulations (Barbosa et al., 2018) and detect genetic variation that boosts fitness in particular environments (Kardos et al., 2021; Razgour et al., 2019). Furthermore, insect species that were once common have been shown to decline quickly (e.g., the rusty-patched bumble bee) (Wagner et al., 2021; Mola et al., 2024; van Klink et al., 2024), highlighting the importance of conducting genetic assessments on common species to inform proactive conservation measures and provide a baseline for future studies (Webster et al., 2023).

### Genetic diversity assessment

We found nucleotide diversity to be comparable to other pierid butterflies (Talla et al., 2019; Zheng et al., 2023) and about 10X lower than the well-studied *Heliconius* butterflies (Martin et al., 2016; Montejo-Kovacevich et al., 2022). We observed slightly higher diversity east of the continental divide, which may reflect differences in historical population size or connectivity between eastern and western populations in this region (Stevens et al., 2012). Higher diversity in eastern populations could also indicate a recent selective sweep (Montejo-Kovacevich et al., 2022) or larger effective population sizes in that region, which may influence adaptive potential (Hare et al., 2011; Hoban et al., 2021). However, additional analyses would be needed to confirm the origins of these patterns, for example explicit tests of demographic history through coalescence simulations and estimating effective population size over time (Allendorf et al., 2021). Above all, this measure of genetic diversity represents a baseline value for which future studies with a greater conservation focus can be compared.

### Evaluating population structure on a regional scale

We used multiple approaches to visualize and quantify population structure for our study area. First, the principal component and *F_ST_* analyses showed that differentiation was relatively weak overall, supporting our prediction of high connectivity throughout the region. Our region-wide multivariate analysis of variance tests showed that differentiation was relatively elevated between eastern and western collection sites. Genetic differentiation across a mountain range has been observed in non-migratory butterflies around the world, for example in a study of altitudinal adaptation in *Heliconius* butterflies in the Andes (Montejo-Kovacevich et al., 2022), and in a closely related species, the orange sulfur (*C. eurytheme*) in the High Sierras in California and Nevada (Dwyer et al., 2015). The marginal signals of east-west differentiation in our study can likely be explained by the presence of the Rocky Mountains and historic biogeography of the region, which has been investigated for a related species (*Colias meadii*) (DeChaine & Martin, 2005). To see if the east-west differentiation is biologically significant for *C. p. eriphyle*, we would need to expand our sampling region across a wider range of latitude.

Additionally, we found patterns of isolation by distance (IBD) across the study region when we regressed geographic distance and genomic distance between sampling sites. IBD was more significant (i.e., slope was steeper) within eastern populations when compared to western populations (Figure S5). IBD is found across butterfly taxa (Lebigre et al., 2022; Trense et al., 2021) and occurs because of landscape features, such as roads, rivers, and forests (Trense et al., 2021) and the quality of the habitat, particularly host plant quality (Lebigre et al., 2022). Additionally, IBD can be generated by the ecology of the butterfly species, with stronger IBD expected in populations with lower dispersal ability, more fragmented or heterogeneous habitats, and narrower host specialization (Lebigre et al., 2022).

Likewise, the patterns observed in our study are likely explained by a combination of the natural history of *C. p. eriphyle* along with landscape characteristics. Clouded sulfurs live in meadows, grasslands, and agricultural fields containing host plants from the Fabaceae (pea) family (Watt et al., 1979). Clouded sulfurs are not particularly strong fliers; dispersal distances are less than 1km on average (Watt et al., 1977) with populations described as sedentary when host plants are at high density, particularly “pest” populations at lower elevations (Tabashnik, 1980). Personal field observations and previous studies have documented the patchiness of meadows in subalpine habitats, with forest stands acting as barriers to dispersal (Keyghobadi et al., 1999; Trense et al., 2021). In the lower elevation areas in the eastern foothills and western slope on either side of the Rocky Mountains, dispersal barriers additionally include roads and housing developments present in these more urban areas (Rochat et al., 2017; Trense et al., 2021). Thus, patterns of genetic differentiation with increased distance are likely driven by the short flight distances and barriers to dispersal present across the sampling region. Additionally, the increased urbanization in eastern Colorado compared to the western slope may help explain why IBD is stronger in the east. Eighty-eight percent of the population of Colorado resides in the Front Range (HRSA, 2022), which comprises the urban centers of Fort Collins, Boulder, Denver, and Colorado Springs. All of the eastern collection sites were located in this region, and the increase in urbanization and human activity likely contributes to the increase in IBD to the east of the divide.

Overall, we suggest that the population structure of *C. p. eriphyle* is comprised of relatively continuous sites, connected via intermediate populations in discrete habitat patches. Dispersal between these intermediate populations generates relatively high gene flow overall, and similarly produce local and regional patterns of isolation by distance and largely panmictic population structure.

### Investigating loci and genes underlying adaptation

Recent work emphasizes that local adaptation across heterogeneous landscapes in populations connected by gene flow (Tigano & Friesen, 2016). In fact, modeling studies show that immigration maximizes local adaptation under heightened environmental variation (Blanquart et al., 2013). For example, gene flow enhanced fitness of wildflower *Clarkia pulchella* under warming conditions (Bontrager & Angert, 2019), and flour beetles adapted to a novel insecticide even with gene flow from a maladapted source population (Durkee et al., 2024).

Similarly, guppies adapted to different levels of predation and maintained phenotypic divergence (bright colors in low-predation habitats, dull colors under high-predation) even with maladapted gene flow (Fitzpatrick et al., 2015). In our system, isolation by distance across the study region suggests substantial connectivity among sites. While gene flow can act as a constraining force to adaptation by swamping adaptive genetic variation (Lenormand, 2002; Slatkin, 1987), gene flow can also facilitate adaptation by maintaining standing variation and introducing adaptive alleles (Carlson et al., 2014; Whiteley et al., 2015). Our study provides evidence that environmentally-associated genetic variation can be maintained despite high gene flow in regional populations of *C. p. eriphyle*, as we identified >16,000 loci located within 25kb of >4,000 genes associated with elevation and summer levels of precipitation and solar radiation.

The results from the genotype-environment association (GEA) analysis strongly suggest that summer precipitation was the leading driver of adaptive divergence among populations, and that this was related to geographic position relative to the continental divide (east vs. west). To further investigate these patterns, we examined the differences in summer precipitation by site and geography (Figure S6). We found significantly greater precipitation (averages of 53mm vs. 39mm) in the eastern sites when compared to the west. We also found greater precipitation at higher vs. lower elevations (averages of 50mm vs. 41mm), but precipitation was higher in the east overall for all sites but two (Figure S6b), highlighting the precipitation differences across the divide. While other correlated variables could also explain the patterns from our GEA, the impacts of precipitation changes on fitness are well studied in Lepidoptera. Summer droughts have been shown to reduce the survival of both larvae and adults (Klockmann & Fischer, 2017), specifically by impacting the availability of and quality of larval food plants and adult nectar sources (Klockmann & Fischer, 2017; van Bergen et al., 2020). For *C. p. eriphyle*, drier summers are associated with increased mortality and decreased abundance in Colorado, where there are 2-5 generations per summer (Watt et al., 1979; Higgins et al., 2015). Thus, we suggest differences in precipitation across the sampled populations, which is supported by historic trends (Schumacher, 2024), largely drives the observed patterns of population adaptation and differentiation across the continental divide in Colorado. Furthermore, higher precipitation in eastern *C. p. eriphyle* populations could lead to decreased mortality and increased abundance in eastern populations, which may underly the pattern of slightly elevated nucleotide diversity that we observed in eastern vs. western sites.

We identified four environmentally-associated loci within the coding regions of genes of interest. First, we investigated genes specifically involved in the production of melanin in butterfly wings, an important trait for thermoregulation (Ellers & Boggs, 2004) and identified two: the *TH* gene, which works to convert tyrosine to dihydroxyphenylalanine (DOPA) at the start of the melanization pathway, and *yellow*, which converts DOPA to melanin (Zhang et al., 2017).

The locus associated with *TH* can be found on the *Z* chromosome (Figure 5b), which has been shown to harbor genetic variation associated with wing coloration in *C. p. eriphyle* (Jahner et al., 2025). Both of these genes had higher allele frequency with greater precipitation, and no significant association with elevation. Numerous past studies have emphasized the importance of melanization for withstanding the selection pressures at increased altitude (Ellers & Boggs, 2002, 2004; Kingsolver & Buckley, 2017; MacLean, Higgins, et al., 2016; MacLean, Kingsolver, et al., 2016). Given that precipitation is positively correlated with altitude (Figure S6), the effects of precipitation and elevation could be somewhat confounded in past studies. Furthermore, it is important to consider evolutionary tradeoffs – butterflies with access to food with greater concentrations of water may be able to allocate more resources into melanin production and adaptation more broadly (Jamieson et al., 2012). Interestingly, the observed increase in melanin-associated allele frequencies with greater precipitation is consistent with Gloger’s rule, which predicts darker pigmentation from increased melanin across taxa in more humid environments (Delhey, 2019). This further suggests that hydric conditions may play a key role in shaping pigmentation-related adaptation in this species, similarly to other insects (e.g., lacewings of the *Chrysoperla* complex) that were mentioned in Delhey’s review (Thierry et al., 2011).

Next, heat shock proteins, including the identified candidate gene *hsp70,* have been well studied in Lepidoptera for their role in thermal stress adaptation (Luo et al., 2015; Mutamiswa et al., 2023). The highest allele frequencies at the SNP located within the coding region of this gene were observed in the drier sites of western Colorado. Remarkably, a study conducted by Higgins et al. in 2015 explored the stress responses of *C. p. eriphyle* populations at two sites in western Colorado, specifically through their expression of *hsp70.* They found that *hsp70* expression was involved in heat stress response, suggesting it may play a role in stress responses more broadly in this species (Higgins et al., 2015).

Finally, cytochromes P450 (including the identified *CYP6B* gene), play a role in detoxification and host plant utilization in caterpillars (Calla et al., 2017). The SNP within the coding region of *CYP6B6* had the highest allele frequencies in eastern, wetter sites, suggesting precipitation may be important for host plant adaptation in this species. Over the past ∼100 years, *C. p. eriphyle* has incorporated alfalfa into their diets and some populations live exclusively on agricultural alfalfa (Tabashnik 1983). While Tabashnik (1983) did not find strong signals of host associated differentiation, whether cytochromes P450 are involved in the switch from native legumes to alfalfa could be explored further in future studies.

Overall, the large number of environmentally-associated loci and the candidate genes involved in key physiological pathways demonstrate that genomic signals of local adaptation can be maintained even with considerable gene flow. Our findings indicate that environmental variation across the continental divide – particularly summer rainfall, which is associated with fitness of *C. p. eriphyle* – underlies the observed adaptative divergence across the region, enabling the species to occupy a broad range of habitats. Taken together, the high levels of both gene flow and adaptive genetic variation should help prevent local expirations and buffer populations against future environmental change.

### Conclusions and implications

For the populations of *C. p. eriphyle* assessed in our study, we observed strong evidence of local adaptation, likely driven by environmental differences east vs. west of the continental divide, despite minimal population structure. We found differences in summer rainfall across the divide drives most patterns of adaptation when compared to elevation and solar radiation. Our exploration of genomic isolation by distance suggests that gene flow likely occurs between low- and higher-elevation populations via intermediate habitat patches, confirming the observations from Watt and colleagues in the 1970s. Due to their low dispersal capabilities, movement across the elevational gradient separating low and high sites likely takes several generations, which would enable rapid adaptation to temperature and other climatic differences present along the gradient (Kingsolver & Buckley, 2017). Our study suggests several genes associated with melanization, thermal tolerance, and detoxification (*TH, yellow, hsp70,* and *CYP6B6*) may underlie adaptation to new environments. Moving forward, monitoring the population genomics of *C. p. eriphyle* and other common Rocky Mountain butterfly species is paramount, particularly given the increases of drought, wildfire, and heat waves in the region (Liu et al., 2015; McCoy et al., 2022), to monitor for declines in genetic health or maladaptation to ongoing and worsening environmental challenges.

The clouded sulfur butterfly has been used as a model system in the past to study insect host evolution and adaptive responses to elevation (Buckley & Kingsolver, 2019; Ellers & Boggs, 2004; Kingsolver, 1983; Kingsolver & Buckley, 2017; Tabashnik, 1983). Here, we add a novel genomic component to the study of *C. p. eriphyle*, suggesting that the genetic basis of local adaptation is strongly associated with summer rainfall, solar radiation, and other environmental variables that vary across elevation and geographic region. These patterns persist despite minimal population structure across the region, highlighting how populations can exhibit patterns of local adaptation even in the face of gene flow. This work provides compelling evidence for how adaptive genomic variation can be maintained amid ongoing gene flow and environmental variability.

## Supporting information

Colias-GEA-supp_mat

## Acknowledgements

This work was supported by the U.S. National Science Foundation Graduate Research Fellowship (award no. 006784) and grants from the Lepidopterists’ Society and the Graduate Degree Program in Ecology at Colorado State University. Thank you to lab and collections managers Christine Rayne and Mackenzie Woods for training and support with molecular biology techniques, and to undergraduate researchers Lindsay Buckentine and McKenna Gonzalez for assistance with data processing. We thank several volunteer field assistants, including Ellen Durkee, Jakub Jarník, and Evi Buckner-Opler, for their support with butterfly collections and field logistics. Field access was provided by Rocky Mountain National Park, the US Forest Service, City of Fort Collins, and numerous private landowners. This work used the RMACC Alpine supercomputer, which is supported by the National Science Foundation (awards ACI-1532235 and ACI-1532236), the University of Colorado Boulder, and Colorado State University. The findings and conclusions in this article are those of the authors and do not necessarily represent the views of the U.S. Fish and Wildlife Service.

## Data Availability Statement

The genomic data that support the findings of this study will be uploaded to Dryad upon acceptance of the manuscript. Scripts are openly available on Github at https://github.com/lilyd-csu/colias-gea/.

## Author Contributions

LFD, KCR, PAO, and RAH conceptualized the study. LFD and RAH acquired funding. LFD, PAO, and RAH collected the samples. LFD completed the lab work with resources and methods support from KCR. LFD, CMB, and BRF performed the genomic analyses and data visualization. LFD wrote the original manuscript draft and all authors (except PAO, who passed away prior to submission) participated in revision.

## Notes

https://github.com/lilyd-csu/colias-gea

### Competing Interest Statement

The authors have declared no competing interest.

