## Supplementary material for "Summer rainfall drives adaptation with gene flow in a widespread butterfly": Colias-GEA-supp_mat

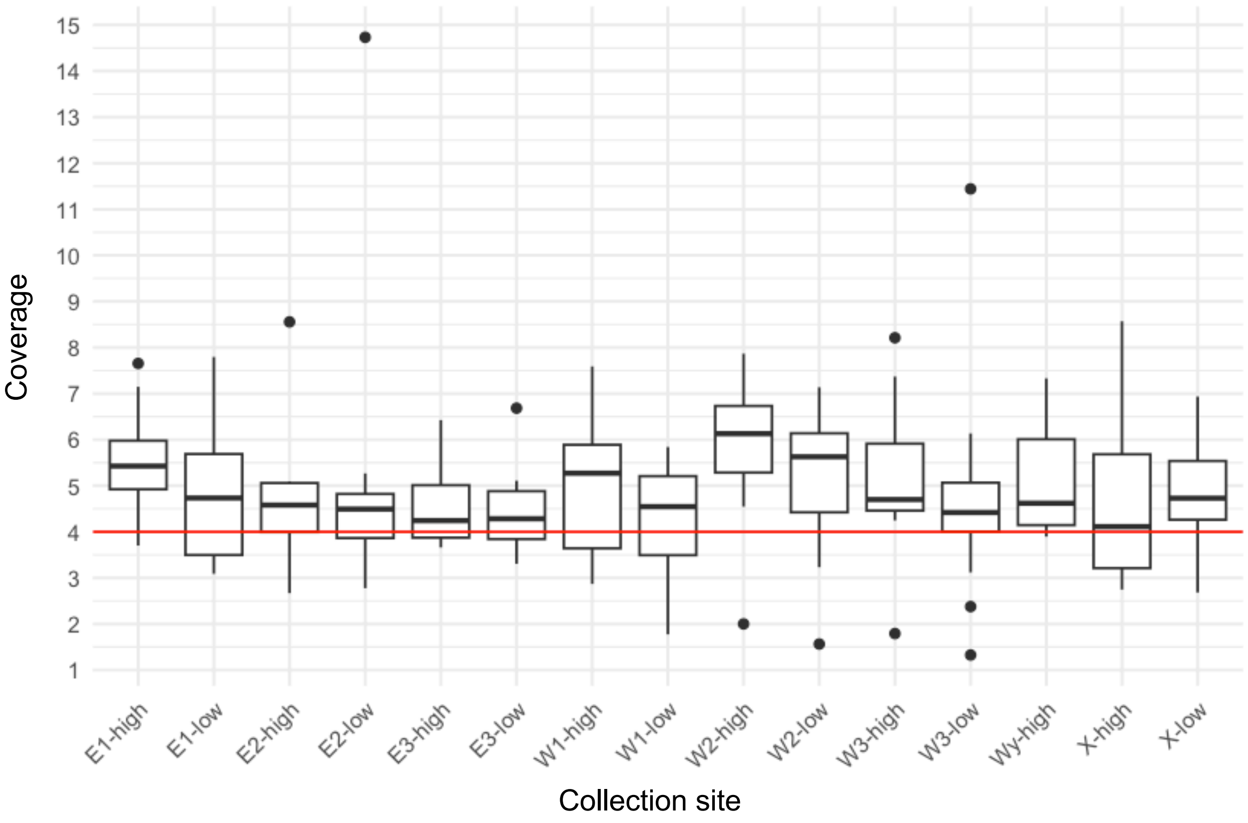


**FIGURE S1. Variation in read depth coverage among collection sites.** Due to the variation in coverage, we downsampled to 4X coverage (red line).

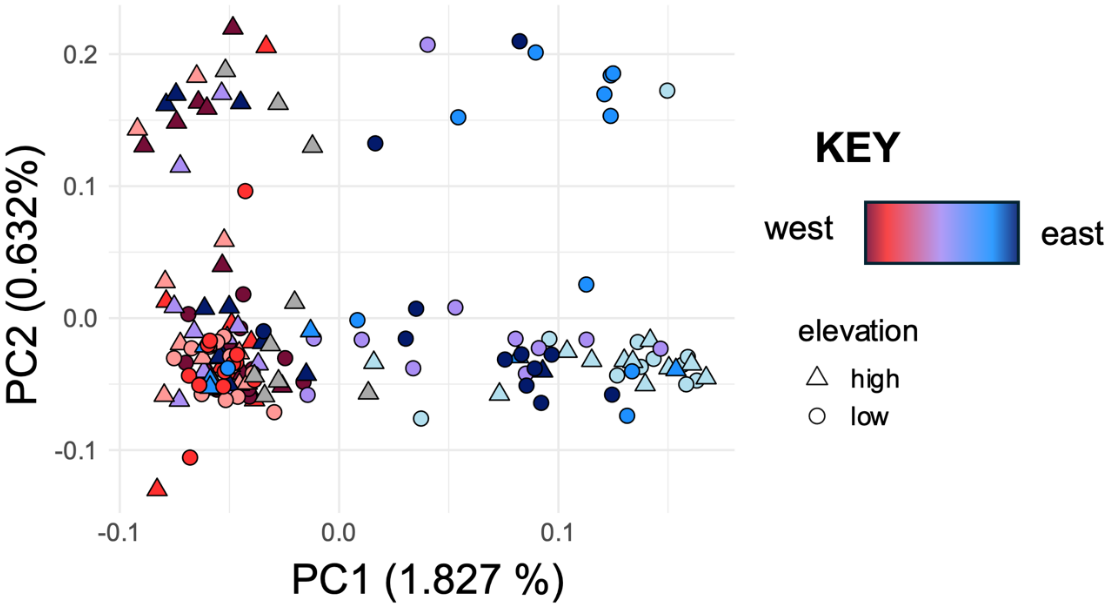


**FIGURE S2. Principal component analysis generated using genotype probabilities** with *pcangsd*. This was used to compare against the analysis done with GATK4*.*

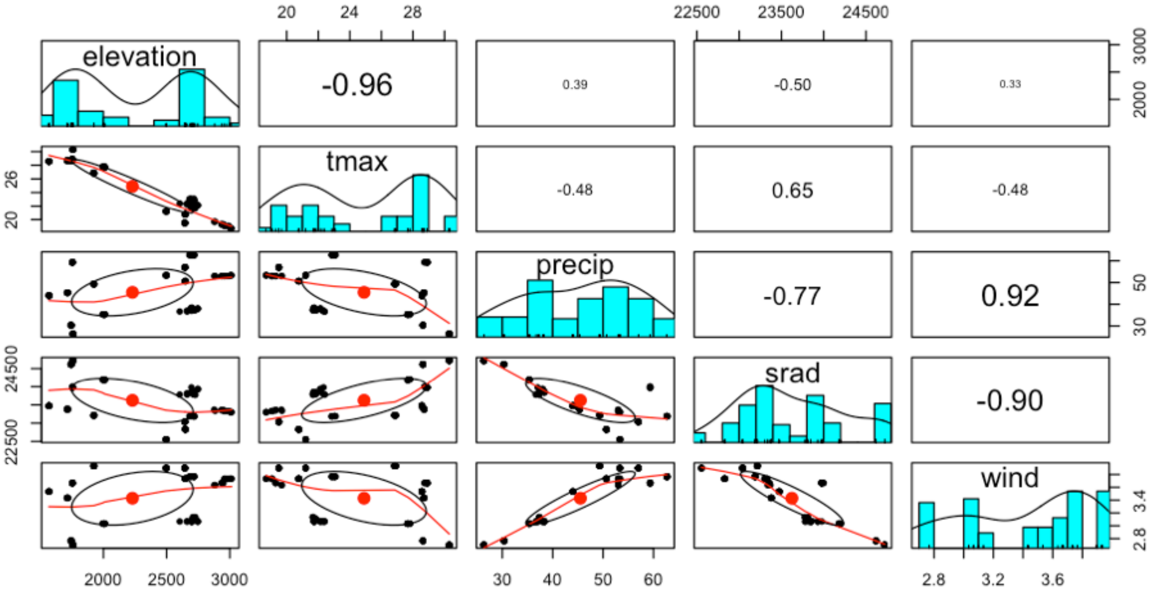


**FIGURE S3. Pearson correlation coefficient (r)** for the environmental variables considered for the redundancy analysis. We included variables with |r| < 0.80, and therefore selected elevation (m), summer monthly precipitation (precip, mm/month), and summer solar radiation (srad, kJ/m$^2$/day). We eliminated max temperature (tmax, °C), which was highly correlated with elevation (*r* = -0.96) and wind (m/s), which was highly correlated with precipitation and solar radiation (*r* = 0.92 and -0.90, respectively).

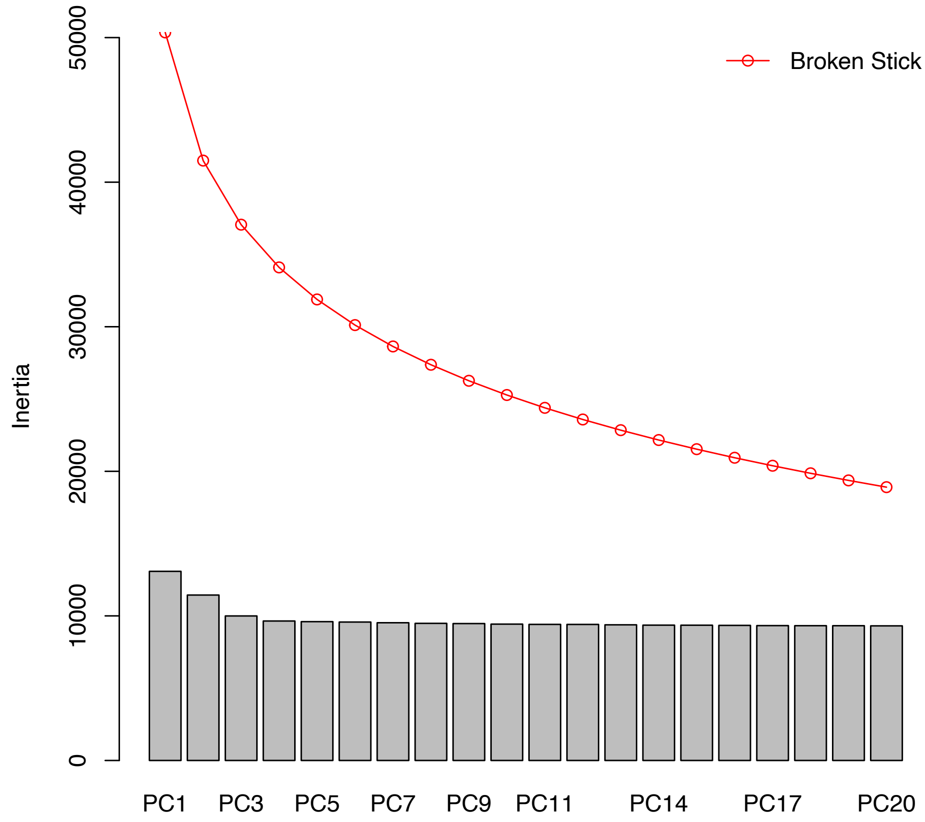


**Figure S4.** Scree plot showing broken-stick criterion for the PCA (Forester et al., 2018).

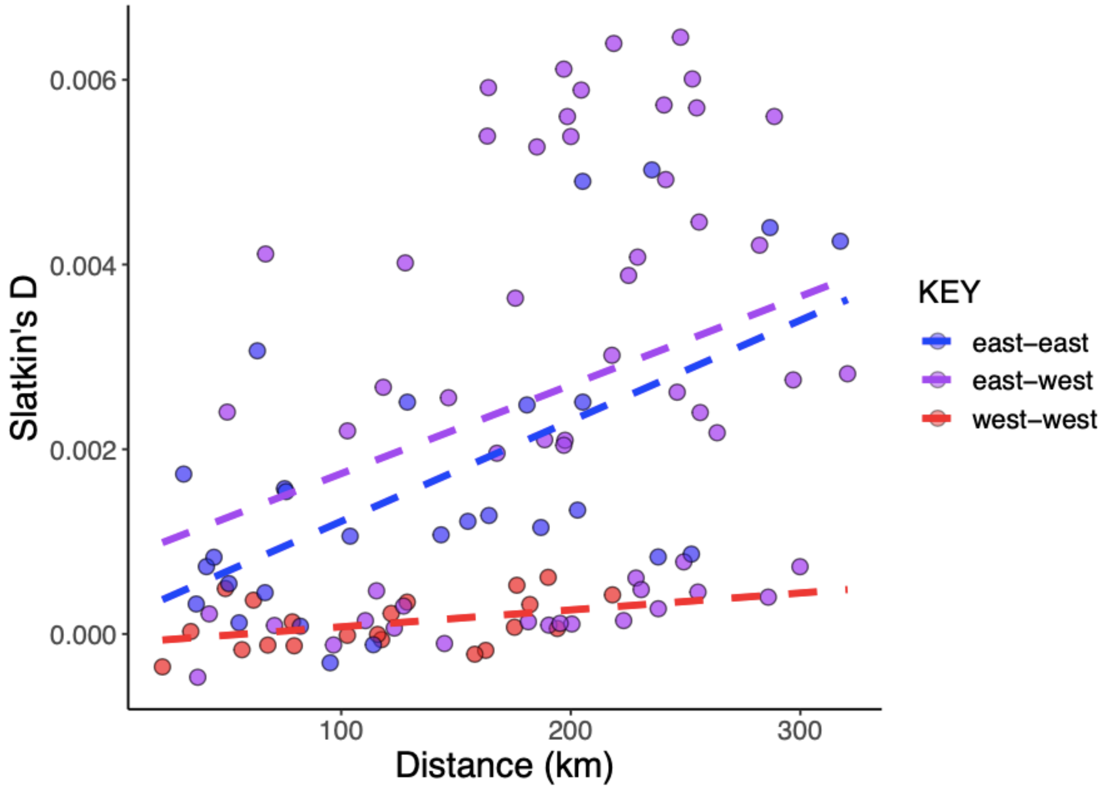


**FIGURE S5. Differences in isolation by distance (IBD)**, color-coded by region. Each point on an IBD plot represents the geographic and genomic distances between two sampling sites. Thus, the key corresponds to whether the comparison is between two sites that are both on the eastern side of the divide ("east-east", blue), both west ("west-west", red), or if one site was east and west ("east-west", purple). The regression lines show the strength of IBD for each group, and show that IBD is stronger (i.e., slope is steeper) for the east-east and east-west comparisons.

(a)

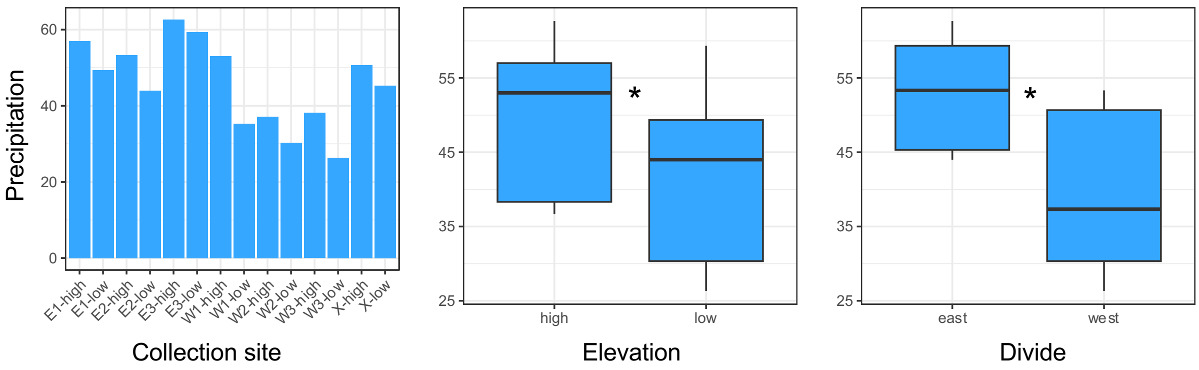


(b)

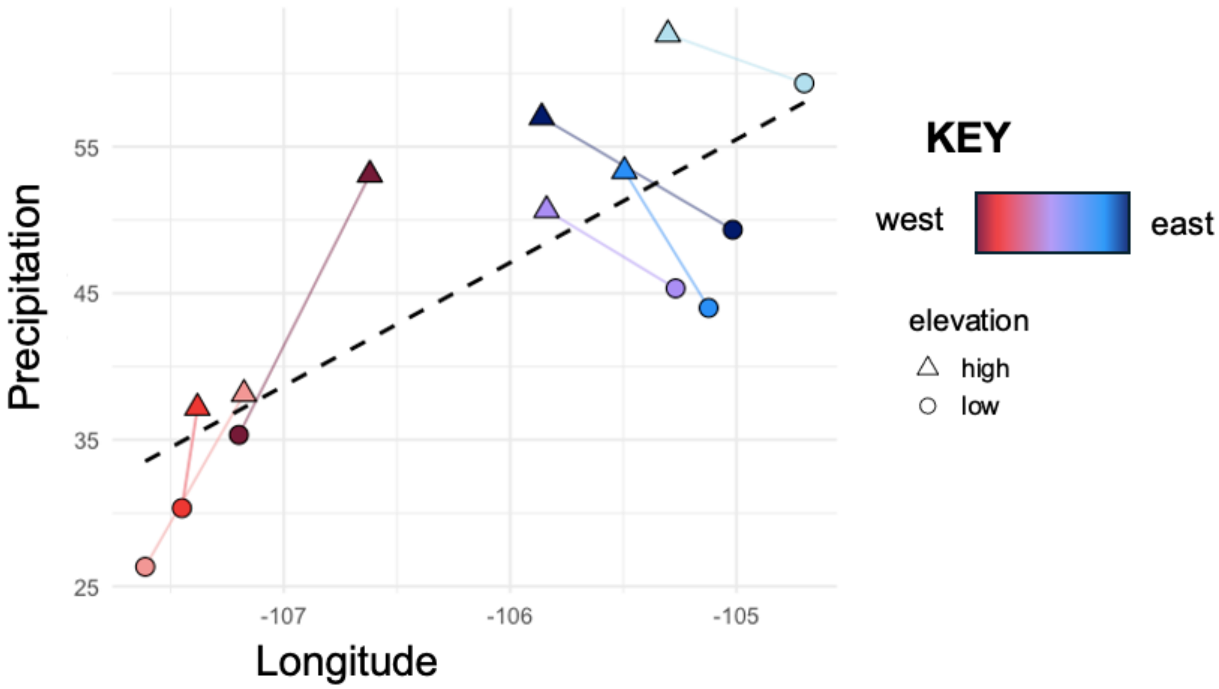


**FIGURE S6. (a) Summary of average monthly precipitation (mm)** included in the redundancy analysis as extracted from *BioClim* for the summer months (June-August) from 1970-2000. Comparisons are shown for all sites, between high (>2,500m) vs. low (<2,000m) sites, and sites east vs. west of the continental divide. Asterisks (*) indicate that the difference in means was significantly greater than zero (t-test, $p < 0.001$). **(b) The correlation of precipitation with longitude**, with lines connecting paired sites. Overall, high-elevation sites (triangles) received higher precipitation than their low-elevation pairs (circles), and eastern sites had higher precipitation than all but two western sites (purple triangle, dark red triangle).
